# Low rate of somatic mutations in a long-lived oak tree

**DOI:** 10.1101/149203

**Authors:** Namrata Sarkar, Emanuel Schmid-Siegert, Christian Iseli, Sandra Calderon, Caroline Gouhier-Darimont, Jacqueline Chrast, Pietro Cattaneo, Frédéric Schütz, Laurent Farinelli, Marco Pagni, Michel Schneider, Jérémie Voumard, Michel Jaboyedoff, Christian Fankhauser, Christian S. Hardtke, Laurent Keller, John R. Pannell, Alexandre Reymond, Marc Robinson-Rechavi, Ioannis Xenarios, Philippe Reymond

**Affiliations:** Center for Integrative Genomics, University of Lausanne, 1015 Lausanne, Switzerland; Department of Ecology and Evolution, University of Lausanne, 1015 Lausanne, Switzerland; Evolutionary Bioinformatics Group, Swiss Institute of Bioinformatics, 1015 Lausanne, Switzerland; Vital-IT Competence Center, Swiss Institute of Bioinformatics, 1015 Lausanne, Switzerland; Department of Plant Molecular Biology, University of Lausanne, 1015 Lausanne, Switzerland; Fasteris SA, 1228 Plan-les-Ouates, Switzerland; Swiss-Prot group, Swiss Institute of Bioinformatics, 1211 Geneva, Switzerland; Risk Analysis Group, Institute of Earth Sciences, University of Lausanne, 1015 Lausanne, Switzerland

## Abstract

Because plants do not possess a proper germline, deleterious somatic mutations can be passed to gametes and a large number of cell divisions separating zygote from gamete formation in long-lived plants may lead to many mutations. We sequenced the genome of two terminal branches of a 234-year-old oak tree and found few fixed somatic single-nucleotide variants (SNVs), whose sequential appearance in the tree could be traced along nested sectors of younger branches. Our data suggest that stem cells of shoot meristems are robustly protected from accumulation of mutations in trees.

---

Accumulation of deleterious mutations is a fundamental parameter in plant ageing and evolution. Because the pedigree of cell division that generates somatic tissue is poorly understood, the number of cell divisions that separate zygote from gamete formation is difficult to estimate; this number is expected to be particularly large in trees and could in theory lead to a large number of DNA replication errors^1–3^. Tree architecture is determined by the modular growth of apical meristems, which contain stem cells. These cells divide and produce progenitor cells that undergo division, elongation and differentiation to form a vegetative shoot, the branch. Axillary meristems are formed at the base of leaf axils and are responsible for the emergence of side branches. They are separated from apical meristems by elongating internodes. In oak, early and repeated growth cessation of terminal apical meristems leads to a branching pattern originating from such axillary meristems. In turn, axillary meristems grow out and produce secondary axillary meristems. This process is reiterated indeterminately to produce highly ramified trees of large stature, resulting in thousands of terminal ramets^4^. The cumulative number of cell divisions separating meristems determines the rate of genetic aging and the potential accumulation of somatic mutations. To detect such mutations, we selected an iconic old oak tree (*Quercus robur*) known as the ‘Napoleon Oak’ by the academic community of the University of Lausanne. The tree was 22 years old when, on May 12, 1800, Napoleon Bonaparte and his troops crossed what is now the Lausanne University campus, on their way to conquer Italy. At the time of sample harvest for our study, the dividing apical meristems of this magnificent tree (Figure 1, Supplementary Figure 1) had been exposed for 234 years to potential environmental mutagens, such as UV and radioactive radiation.

**Figure 1.**
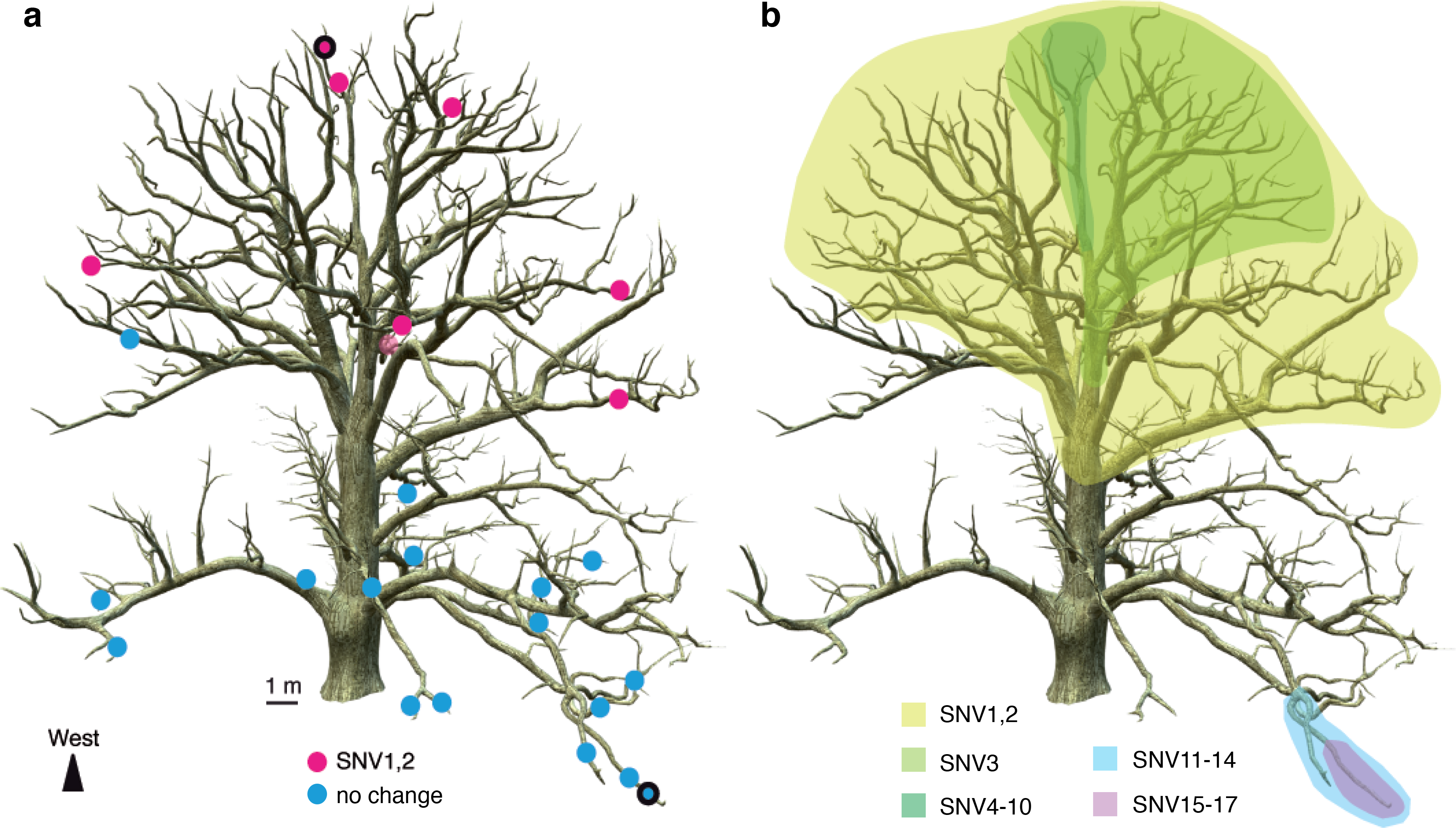
Distribution of somatic mutations in the Napoleon oak. **a**, The genome of two leaf samples utlined dots) was sequenced to identify single-nucleotide variants (SNV). 17 SNVs were confirmed and analysed in 26 other leaf samples to map their origin. A reconstructed image of the Napoleon Oak shows similar ation of two SNVs (magenta dots) on the tree. Blue dots represent genotypes that are non-mutant for these Vs. Three non-mutant samples are not visible on this projection. Location of other SNVs can be found in Supplementary Figure 2. **b**, Location of all identified SNVs. Sectors of the tree containing each group of SNVs represented by different colours.

To identify fixed somatic variants (i.e., those present in an entire sector of the Napoleon Oak) and to reconstruct their origin and distribution among branches, we collected 26 leaf samples from different locations on the tree. We first sequenced the genome from leaves sampled on terminal ramets of one lower and one upper branch of the tree. We then used a combination of short-read Illumina and single-molecule real-time (SMRT, Pacific Biosciences) sequencing to generate a *de novo* assembly of the oak genome. After removing contigs <1000bp, we established a draft sequence of ca. 720 megabases (Mb) at a coverage of ca. 70X, with 85,557 scaffolds and a N50 length of 17,014. Our sequence is thus in broad agreement with the published estimated genome size of 740 Mbp^5^. The oak genome is predicted to encode 49,444 predicted protein-coding loci (Supplementary Table 1).

We used two approaches to identify SNVs (single-nucleotide variants) between the sequenced genomes of the two terminal branches. First, we aligned Illumina paired-reads on the repeat-masked genome in combination with the GATK^6^ variant caller. This allowed us to establish a list of 3,488 potential SNV candidates with a high confidence score. From this list, 1,536 SNVs were experimentally tested by PCR-seq and only seven could be confirmed (see Methods). Second, we used fetchGWI^7^ to map read pairs to the non-masked genome. We were able to call 5,330 potential SNVs from the mapped reads using a simple read pileup process. Further analysis identified 82 putatively variable positions, including the seven already identified using the repeat-masked genome analysis described above (see Methods). Ten of the remaining 75 candidates from the second approach were confirmed by PCR-seq, increasing the total number of confirmed SNVs separating the two genomes to 17 (Figure 1, Supplementary Table 2), which were further confirmed by Sanger sequencing. Based on a conservative estimate, we are likely to have missed no more than 18 further such sites (17 candidates and 1 false negative, see Supplementary Methods).

All 17 confirmed SNVs were heterozygous, as expected for novel somatic mutations. Intriguingly, two SNVs were found on the same contig, separated by only 12 bp (Supplementary Table 2). Sixteen SNVs occurred in introns or non-coding sequences that are probably neutral. The remaining one (SNV1), which occurred in a large sector of the tree, generates an arginine-to-glycine conversion in a putative E3-ubiquitin ligase (Supplementary Table 2). The functional impact of exchanging a positively charged arginine with a non-charged and smaller glycine residue is unknown and deserves further analysis.

Having confidently established 17 SNVs, we then assessed their occurrence throughout the tree. We used Sanger sequencing to genotype the remaining 24 terminal branches sampled from other parts of the tree and checked for the presence of each SNV. SNVs were found in different sectors of the tree in a nested hierarchy that clearly indicates the accumulation of mutations along branches during development (Figure 1, Supplementary Figure 2). These results both provide independent confirmation of the originally identified SNVs, and demonstrate their gradual, nested appearance and fixation in developmentally connected branches during growth of the oak tree. Thus, while the exact ontogeny of the Napoleon Oak may be difficult to reconstruct, our SNV analysis generated a nested set of lineages supported by derived mutations, analogous to a phylogenetic tree.

The spontaneous mutation rate in plants has been estimated to range from 5 × 10^-9^ to 30 × 10^-9^ substitutions/site/generation, based on mutations accumulated during divergence between monocots and dicots^8^. Values for mutation accumulation lines of *Arabidopsis thaliana* maintained in the laboratory range between 7.0 and 7.4 × 10^-9^, which corresponds to ~1 mutation/genome/generation^9,10^. *Arabidopsis* is an annual plant that reaches approximately 30 cm in height before producing seeds. In contrast, the physical distance traced along branches between the terminal branches we sequenced for the Napoleon Oak is about 40 m (Figure 1). Assuming similar cell sizes between oak and *Arabidopsis*, there should have been about 133 times more mitotic divisions separating the extremes of somatic lineages in Napoleon Oak than in *Arabidopsis* (4,000 cm/ 30 cm). Under the assumption that the per-generation mutation rate is correlated with the number of mitotic divisions from zygote to gametes of the next generation^1,11^, and given the ~6-fold larger oak genome, we thus initially estimated that the number of mutations/genome/generation rate should be ~800-fold higher (133 × 6) for the Napoleon Oak than for Arabidopsis, i.e. ~800 SNVs/Napoleon Oak genome. This value is much higher that the 17 SNVs we actually found, even under the conservative assumption that we missed 18 other SNVs from the list of potential variants. Our original hypothesis is thus not supported by the data and another mechanism has to be invoked.

Classical studies of shoot apical meristem organization have reported that the most distal zone has a significantly lower rate of cell division than more basal regions of the apex, and might therefore be relatively protected from replication errors^12,13^. In a recent study that followed the fate of dividing cells in the apical meristems of *Arabidopsis* and tomato, Burian et al.^14^ showed that an unexpectedly low number of divisions separate apical from axillary meristems. In these herbaceous plants, axillary meristems are separated from apical meristem stem cells by seven to nine cell divisions, with internode growth occurring through the division of cells behind the meristem. The number of cell divisions between early embryonic stem cells and terminal meristems thus depends more on the number of branching events than on absolute plant size. Burian et al.^14^ postulated that if the same growth pattern described above for *Arabidopsis* and tomato applies to trees, their somatic mutation rate might be much lower than is commonly thought, and the majority of fixed mutations should be found in relatively small sectors as nested sets of mutations. Napoleon Oak’s apical meristems are of similar diameters to those of tomato^14^ (Supplementary Figure 3) and show similar ontogeny. It thus seems reasonable to suppose that the growth pattern described in *Arabidopsis* and tomato is quite general in flowering plants and might also apply to long-lived trees. The low number of SNVs and their nested appearance in sectors of the Napoelon Oak are thus consistent with hypotheses proposed in Burian et al.^14^.

Mutations accumulate with age, irrespective of plant stature, and long-term exposure to UV radiation contributes to such changes. For the Napoleon Oak, the type of observed SNVs were mostly G:C →A:T transitions, indicative of UV-induced mutagenesis (see Supplementary Discussion). However, oaks protect their meristems in buds under multi-layered leaf-like structures (Supplementary Figure 3), potentially reducing the incidence of UV mutagenesis. The relatively low somatic mutation rate implied by our data may thus be explained by the protective nature of oak bud morphology as well as by the pattern of cell division predicted by Burian et al.^14^. Our results also suggest that mutations due to replication errors in long-lived plants may be less important than environmentally induced mutations. In this context, it is noteworthy that there was no evidence for an expansion of DNA-repair genes in the oak genome compared to *Arabidopsis* (Supplementary Table 3).

To our knowledge, only two examples of functional mosaicism have been reported in trees, a low incidence that might be attributable to the low somatic mutation rate that we report. Although most non-neutral mutations should be maladaptive, eucalyptus trees have been observed with a few branches that are biochemically distinct from the rest of the canopy and have become resistant to Christmas beetle defoliation^15,16^. Functionally relevant somatic mutations, such as SNV1 in our study, may thus occasionally contribute to adaptive evolution if transferred to the fruits, but will more typically increase the genetic load of a population, with implications for inbreeding depression and mating-system evolution (Supplementary Discussion).

Our data give an unprecedented view on the limited role played by somatic mutations in a long-lived organism, and support the view that stem cells in trees, although vulnerable to environment-induced and replication-induced mutations, are probably quite well protected from them. Consistent with this finding, a recent study in *Arabidopsis* has shown that the number of cell divisions from germination to gametogenesis is independent of life span and vegetative growth^17^. Future studies on different tree species and older specimens are needed to test the generality of our study. This work also illustrates the potential for analyses of multiple genomes from single individuals, which throw exciting new light on the rate, distribution and potential impact of somatic mutations in both plant and animal tissues^18,19^.

## Methods

### Materials and genome sequencing

Leaves were collected in April 2012 from the terminal part of a lower (sample 0) and an upper branch (sample 66) of the Napoleon Oak (*Q. robur*) on the Lausanne University Campus (Switzerland, 46°31′18.9″N 6°34′44.5″E). The age of the tree was estimated by a tree ring analysis from a sample taken at the basis of the trunk (Laboratoire Romand de Dendrochronologie, 1510 Moudon, Switzerland). DNA from the two samples was extracted and the genome sequenced. Paired-end sequencing libraries with insert size of 400 bp were constructed for each DNA sample according to the manufacturer’s instructions. Then, 100 bp paired-reads were generated on Illumina HiSEq 2000 at Fasteris (www.fasteris.com). In addition, 3 kb mate-pair libraries from sample 0 were constructed and sequenced with single-molecule real-time (SMRT) technology according to the manufacturer’s instructions (Pacific Biosciences). Short reads were combined with PacBio reads to assemble a reference genome (Supplementary Methods).

### SNV identification

We used two different methods to identify SNVs (see flowchart, Supplementary Figure 4). In the first one, Illumina reads (278,547,120 and 278,651,792 for sample 0 and 66, respectively) were aligned to the masked (RepeatMasker, v4.05) *de novo* assembly with Bowtie2 (v2.2.2, https://sourceforge.net/projects/bowtie-bio/files/bowtie2/2.2.2) using default parameters. GATK^6^ v2.5.2 was used for local realignment and variant calling using standard hard filtering parameters according to GATK Best Practices recommendations^20^. Prior to variant calling, each sample was screened for duplicates using PICARD tools (http://broadinstitute.github.io/picard/v2.9.0, MarkDuplicates). Variants with confidence score ≥50 were retained further. We identified 1,832,554 heterozygous sites common to both samples, as well as 314,865 putative differences between sample 0 and 66 (165,489 sites predicted to be homozygous on sample 0 and heterozygous on sample 66 and 149,376 homozygous on sample 66 and heterozygous on sample 0). The distribution of the confidence scores of the 1,832,554 heterozygous sites common to both samples was a superposition of a Gaussian distribution, peaking at 910, possibly representing true positives, and of an exponential distribution, possibly representing the decreasing number of false positives with regard to increasing confidence score. Importantly, the distribution of scores of the sites with putative differences between samples was an exponential distribution of very low values, similar to the potential false positives of shared heterozygote sites. We thus hypothesized that sites that are truly different between samples 0 and 66 were unlikely to be present at sites with a confidence score below 300. From 3,488 putative SNVs with a confidence score ≥300 on the heterozygous sites and ≥200 on homozygous sites, we selected 1,536 SNVs for validation by PCR-seq (Supplementary Methods). We identified only 7 true SNVs that were further confirmed by Sanger sequencing. This low rate is consistent with the expectation from the distribution of GATK scores for these sites.

In the second method, Illumina reads of samples 0 and 66 were mapped against the non-masked oak genome assembly. The genome was 719,779,348 bp long, but 69,130,634 (9.52%) of those nucleotides were gaps and were discarded, leaving an actual search space of 650,648,714 bp. Of the latter, 458,143,725 nucleotides with a read coverage ≥8 in both samples were analysed further. The mapping process was performed at the read pair level by the genome mapping tool, fetchGWI^7^, followed by a detailed sequence alignment tool, align0^21^. Potential SNVs were called from the mapped reads by a simple read pileup process followed by detection of positions where the pileup shows variations with respect to the reference genome; this produced a list of 5,330 positions. Those positions were browsed through a local adaptation of the samtools pileup browser^22^ to evaluate the quality of the mapping in the surrounding region and to discriminate between well-assembled high-quality regions with two alleles per sample, or low complexity and possibly badly assembled repeated regions. Criteria for selection were ≥8 reads in each orientation (see above); 100% homozygosity site for one sample and at least 30% minor allele frequency for the other sample with variants in both orientations; and coherent sequence ±50 bp from variant site. This manual process led to the selection of 82 putative variable positions, including the seven already identified. Upon experimental validation, 10 of the remaining 75 candidates were confirmed by PCR-seq and Sanger sequencing. The Food and Drug Administration (FDA) has evaluated this approach in an effort to assess, compare, and improve techniques used in DNA testing on human genome variation analysis (https://precision.fda.gov/challenges/consistency). Within this frame, our method reached a F-score (F-score evaluates precision and recall) over 95% comparable to other identifiers like BWA coupled with GATK.

### SNV Genotyping

Leaf DNA from different locations on the tree was prepared and amplified using primers located 100-150 bp away from the 17 confirmed SNVs (Supplementary Table 4). Amplicons were then subjected to Sanger sequencing.

### Data availability

All Illumina reads and SMRT sequences have been deposited in GenBank under accession BioProject PRJNA327502.

## Acknowledgements

This work was funded by the University of Lausanne through a supportive grant from the University rectorate and by the Swiss National Science Foundation (Agora Grant CRAGI3_145652). The Pacific Biosciences RS II sequencing was performed at the Lausanne Genomic Technologies Facility (GTF). The purchase of the GTF’s RS II instrument was financed in part by the *Loterie Romande* through the *Fondation pour la Recherche en Médecine Génétique.* We thank Keith Harshman, Johann Weber, and Mélanie Dupasquier from the GTF for sequencing. We thank Cris Kuhlemeier for sharing unpublished results, Jean Tercier for tree-ring analysis, Woodtli+Leuba SA for sample collection, Nicolas Guex for advice on SNV calling and Jean-Jacques Strahm and Marco Bonetti for providing oak images.

## Author contributions

F. sequenced the genome. E.S.-S., S.C., M.P. assembled and annotated the genome. N.S., E.S.-S., C.I. identified SNVs. C.G.-D., J.C. extracted DNA and confirmed SNVs. E.S.-S., M.R.-R. analyzed genome duplication. P.C. produced cross-sections of oak apical meristems. S. established a list of DNA repair genes. F. S. provided statistical help with the analyses. J.V., M.J. produced a 3D model of the oak tree. C.H., C.F., L.K., I.X., M.R.-R., J.P., A.R., P.R. conceived the project and wrote the manuscript.

## Additional information

Supplementary information is available for this paper.

Correspondence and requests for materials should be addressed to P. R.

---

**Supplementary Figure 1.**
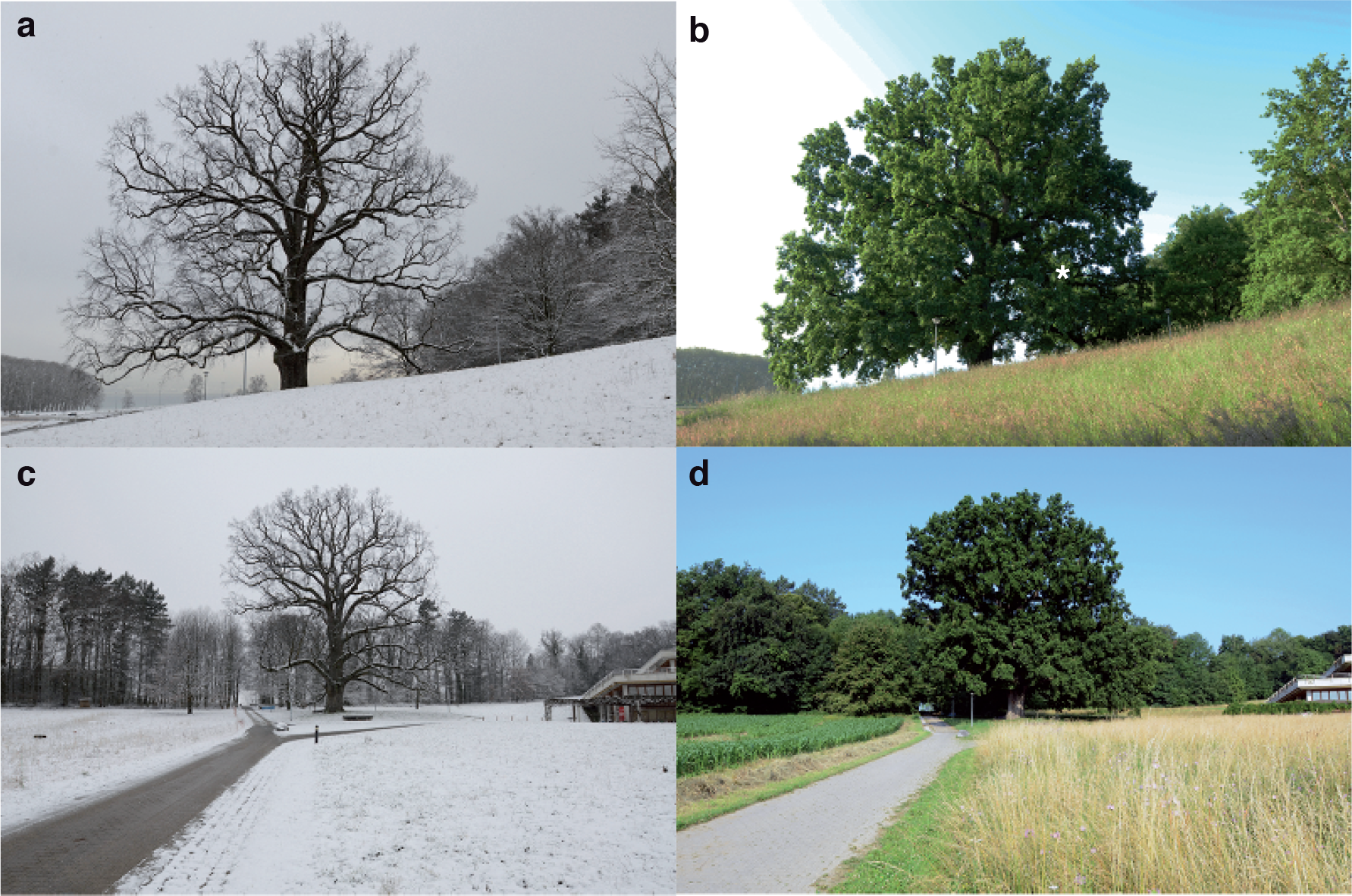
Napoleon Oak. Photographs of the Napoleon Oak on the Lausanne University campus taken in winter and summer. **a, b**, South view. **c, d**, North-West view.

**Supplementary Figure 2.**
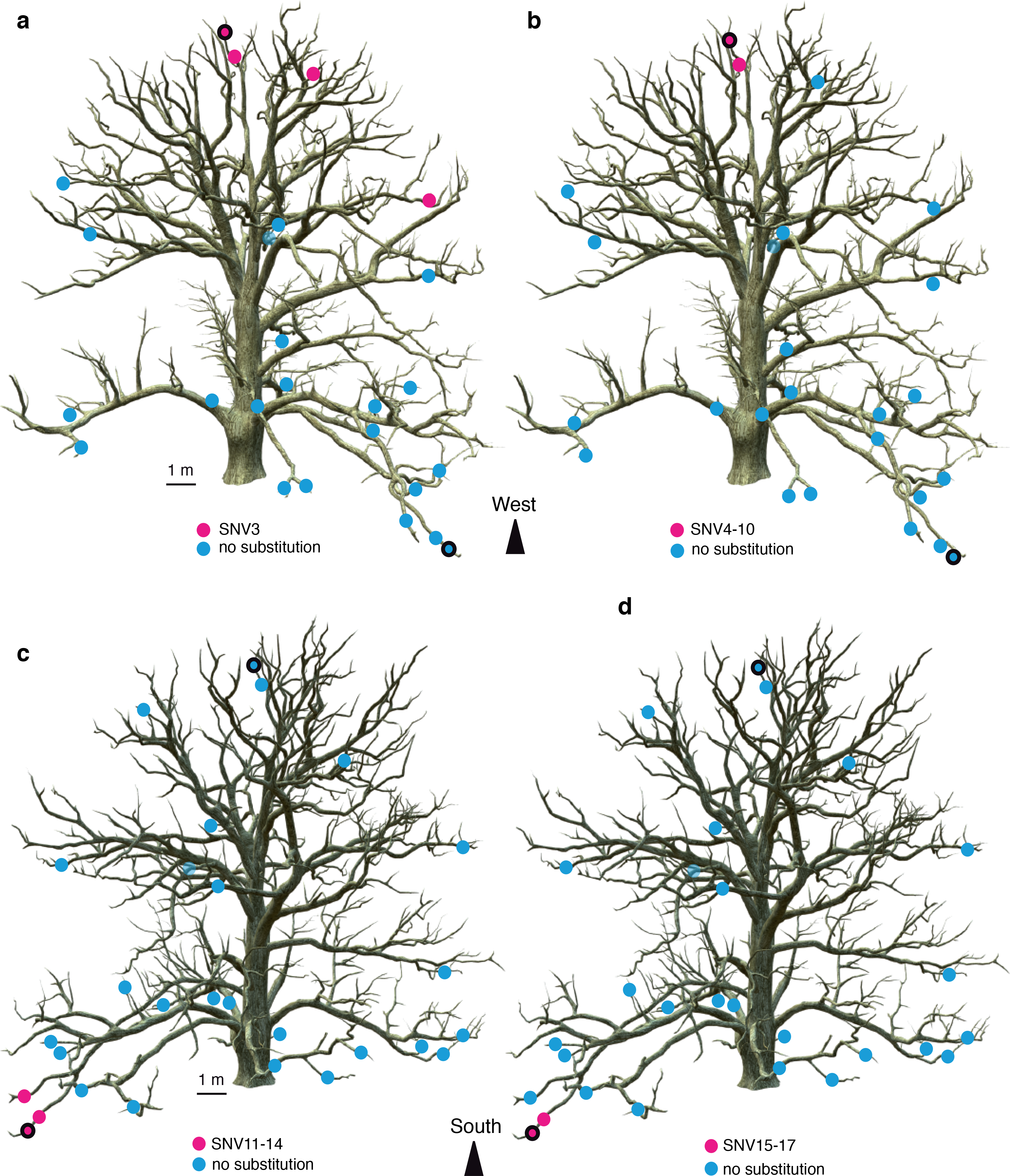
Distribution of somatic mutations in the Napoleon Oak. The genome of two leaf samples (outlined dots) was sequenced to identify single-nucleotide variants (SNV). 17 SNVs were confirmed and analysed in 26 other leaf samples to map their origin. **a-d**, Reconstructed images of the Napoleon Oak show the location of different SNVs (magenta dots) on the tree. Blue dots represent genotypes that are non-mutant for these SNVs.

**Supplementary Figure 3.**
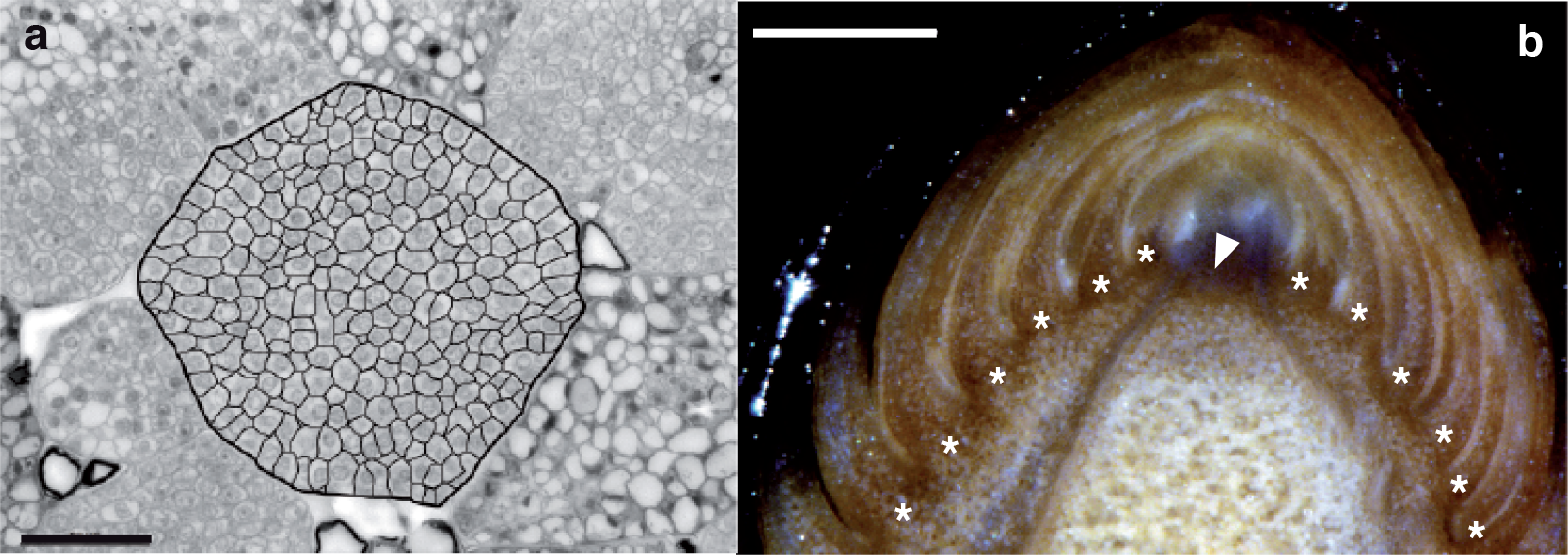
Napoleon Oak apical meristem. **a**, Cross-section of an apical meristem. Meristematic cells are delineated. Surrounding cells belong to leaf-like structures surrounding the meristem. Scale bar, 50 mm. **b**, Longitudinal section of an apical bud. Apical meristem (arrowhead) is surrounded by leaf-like structures (stars). Scale bar, 500 mm.

**Supplementary Figure 4.**
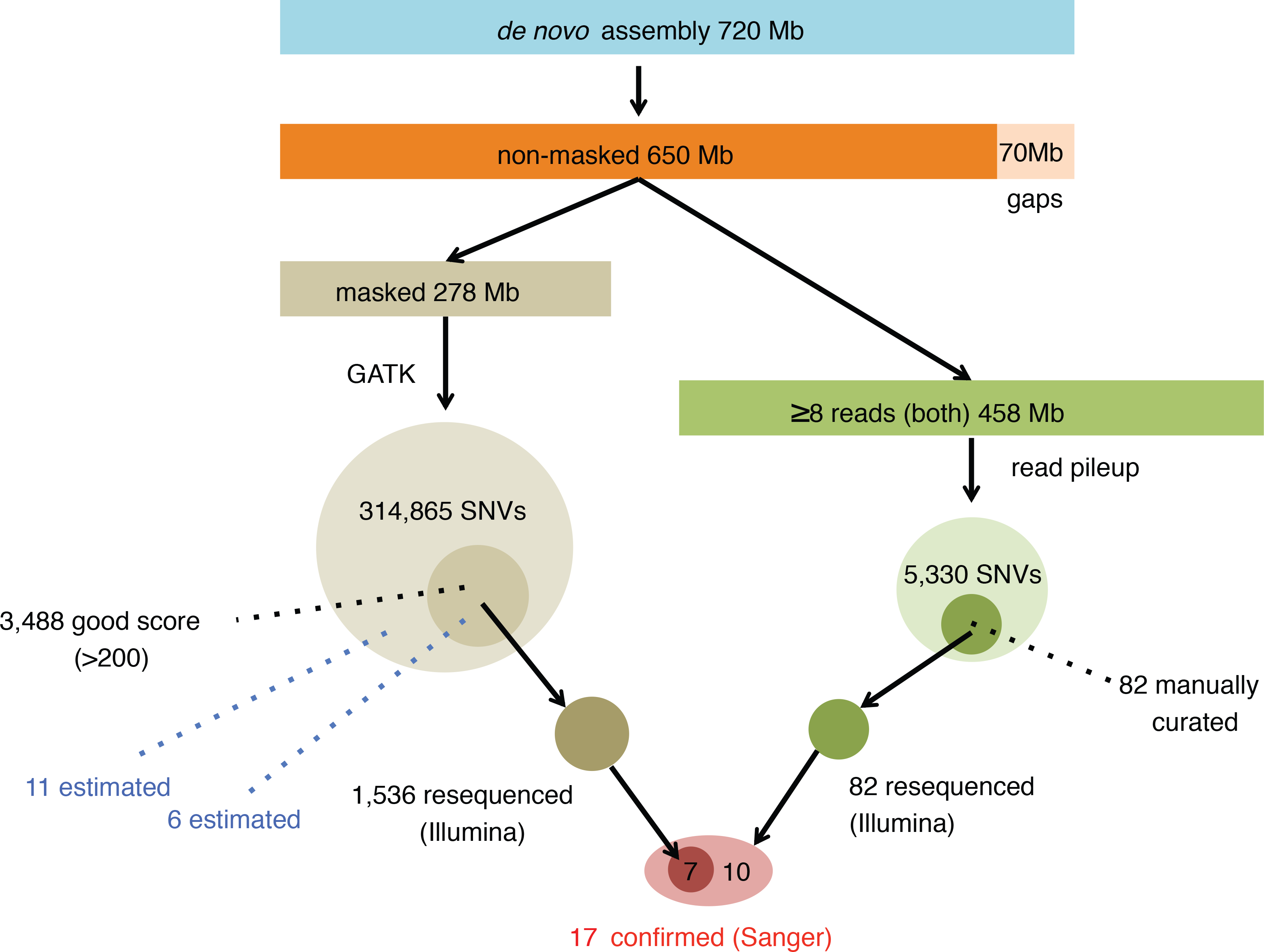
Flowchart of SNV identification methods.

**Supplementary Figure 5.**
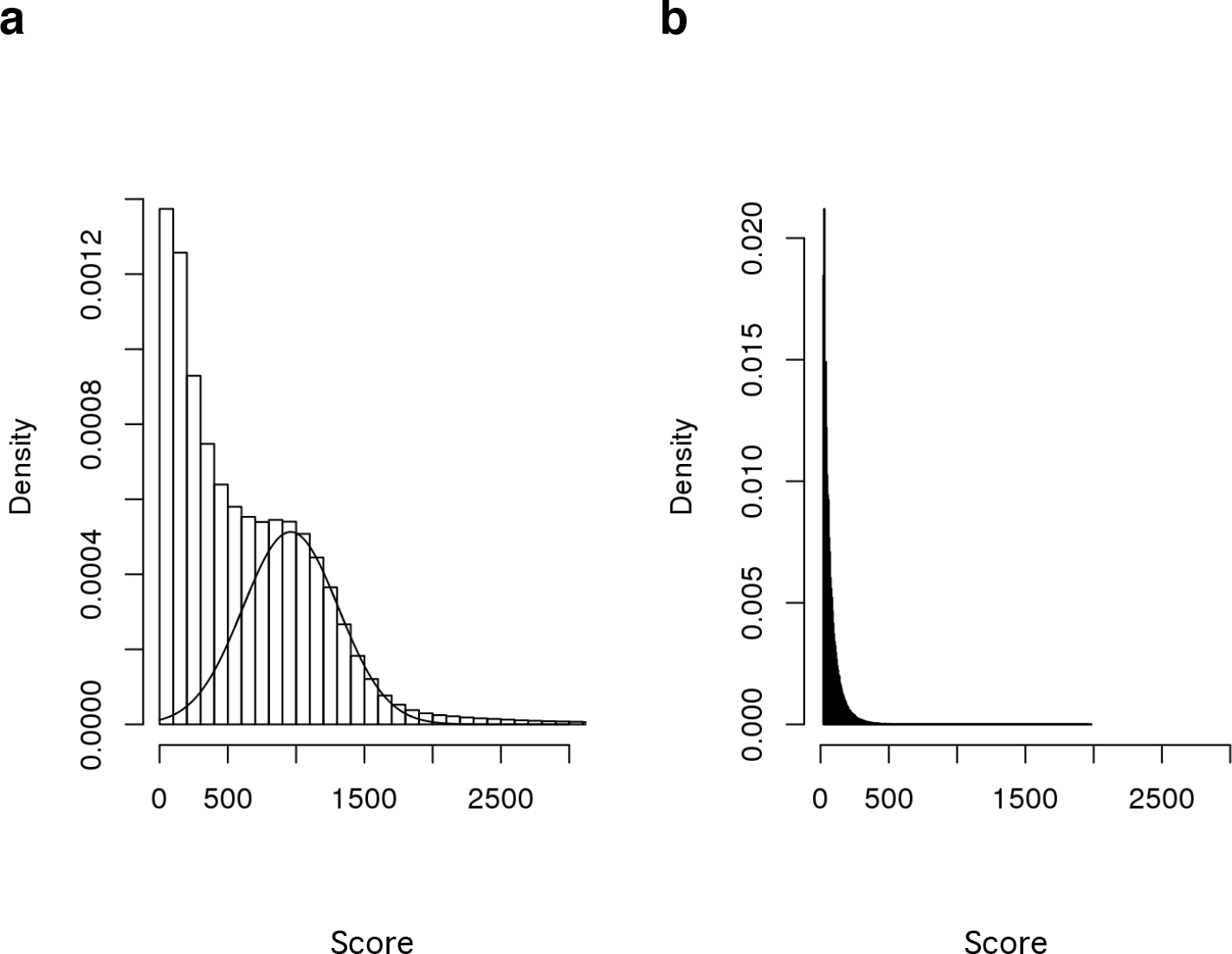
Distribution of variants. **a**, Distribution of the confidence scores of the 1′832′554 heterozygous sites common to both samples 0 and 66. **b**, Distribution of the confidence scores of 165′489 heterozygous sites in sample 66 that are homozygous in sample 0.

**Supplementary Figure 6.**
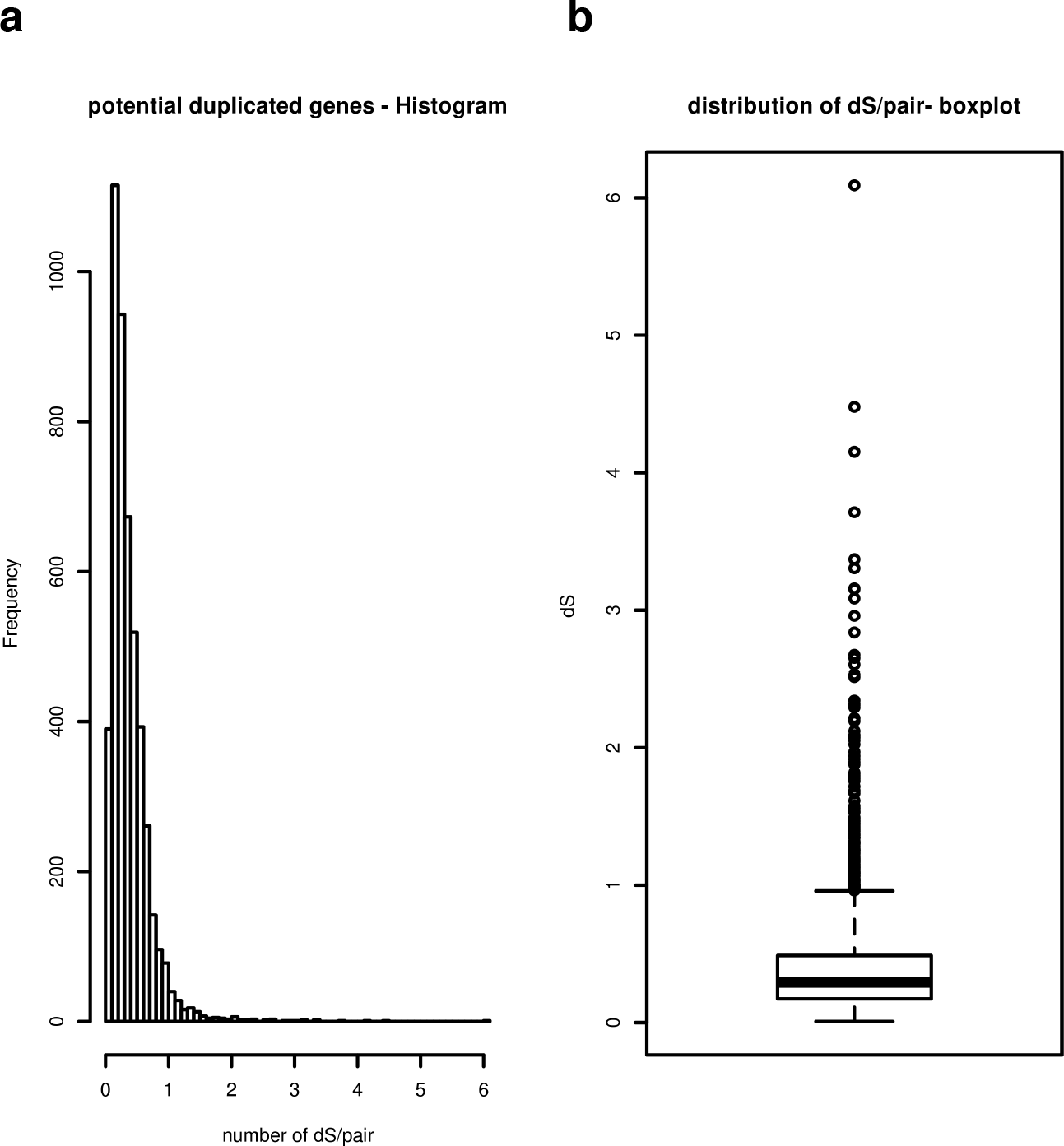
Analysis of oak genome duplication. **a**, Frequency plot and **b**, box plot showing the distribution of synonymous distances (dS) on a stringent set of 4,777 paralog pairs. This analysis was done with a threshold BLAST E-value <1e-10 and by removing multigene families of more than 20 members.

**Supplementary Figure 7.**
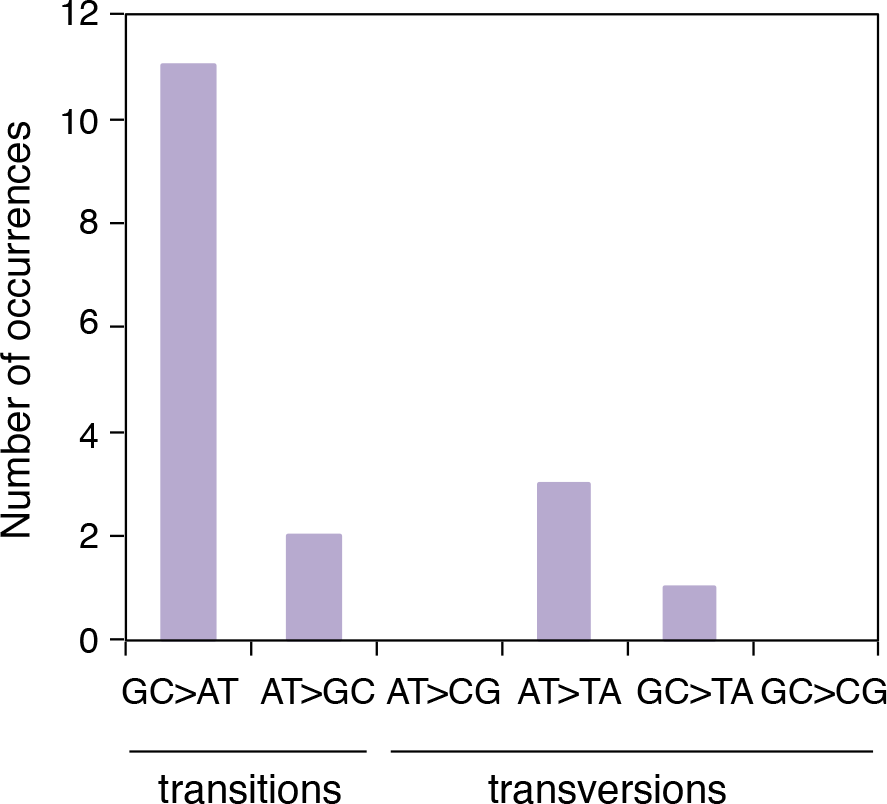
Spectrum of somatic mutations between two Napoleon Oak genomes. The type of substitution for 17 confirmed oak SNVs is shown.

**Supplementary Table 1:** *Quercus robur* genome statistics.

|  |  |
| --- | --- |
| **Genome** |  |
| Total genome length (bp) | 719,779,348 |
| Number of scaffolds | 85,557 |
| Maximum scaffold length (bp) | 317,245 |
| NG50 based on 740 Mbp (bp) | 17,014 |
| Gaps (%) | 9.52 |
| Masked (%) | 39.84 |
| **Genes** |  |
| Average length (bp) | 2,360 |
| Maximum length (bp) | 47,221 |
| Average intron length (bp) | 740 |
| Average exon length (bp) | 232 |
| **Proteome** |  |
| Total predicted proteins | 49,444 |
| Full proteins | 44,096 |
| Partial proteins | 5,348 |
| Nb proteins with orthologous in *Glycine max* | 39,656 |
| Nb orthologous in *Glycine max* + functional annotation | 16,323 |
| Nb orthologous in *Glycine max* + function via ATH | 23,333 |

**Supplementary Table 2.**
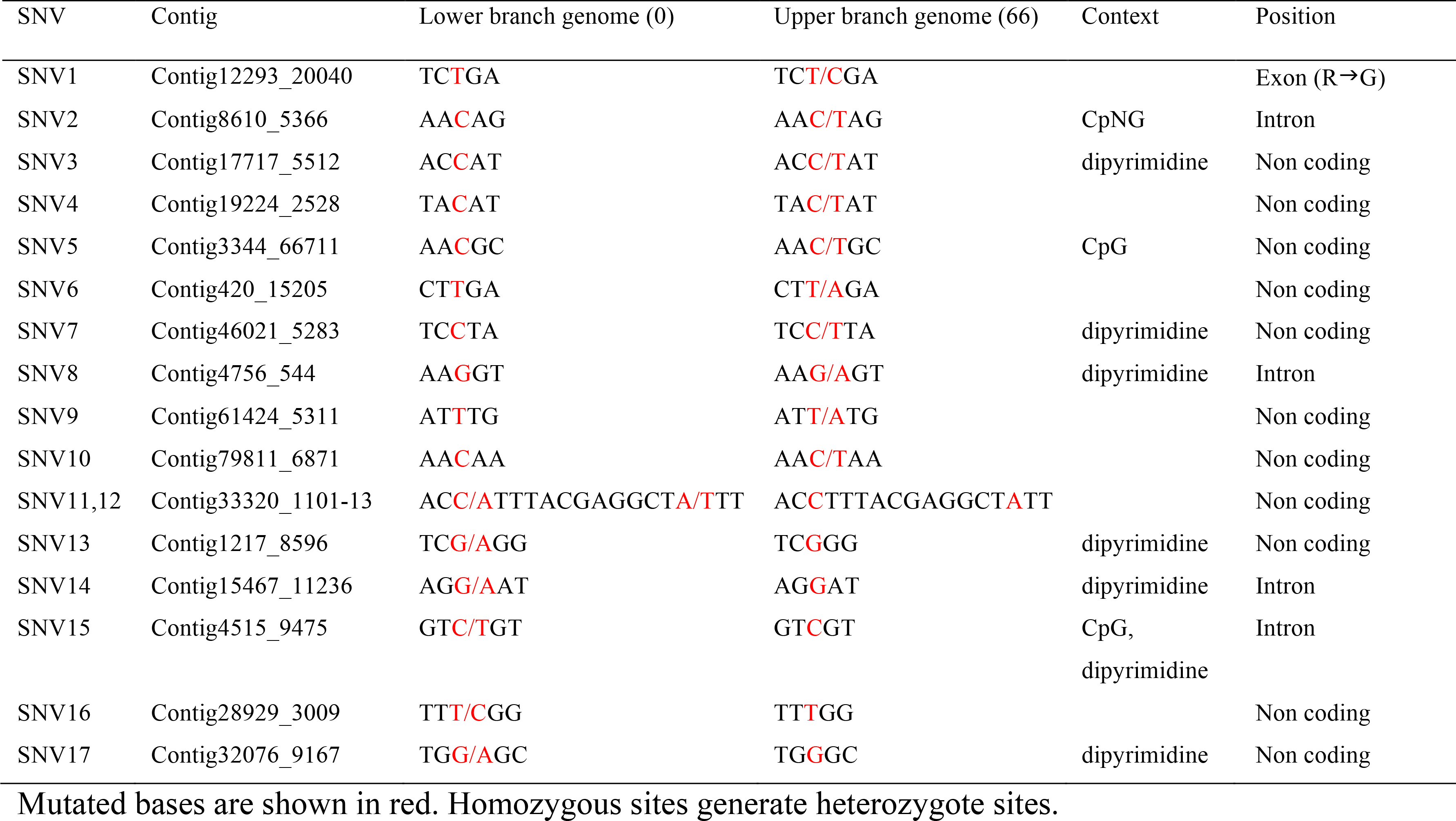
SNVs in the Napoleon Oak.

**Supplementary Table 3.** Orthology and duplication of DNA repair genes in oak and peach trees.

| Branch of the phylogeny | DNA repair genes |  |  | All genes |  |  |
| --- | --- | --- | --- | --- | --- | --- |
|  | duplicated | total | % total | duplicated | total | % total |
| *Quercus robur* | 0 | 228 | 0.0 | 258 | 10,199 | 3.5 |
| *Prunus persica* | 5 | 228 | 2.2 | 860 | 16,004 | 5.4 |
| *Quercus* - *Prunus*<br>common ancestor | 1 | 228 | 0.4 | 523 | 8,474 | 6.2 |

All gene counts are for *Arabidopsis thaliana* orthologs; the number of "all genes" varies according to the number of orthologs detected for each set (i.e., from *Quercus robur*, from *Prunus persica*, or shared). All orthology detection and lineage-specific duplication calls are from OMA (see Experimental Procedures)

**Supplementary Table 4.**
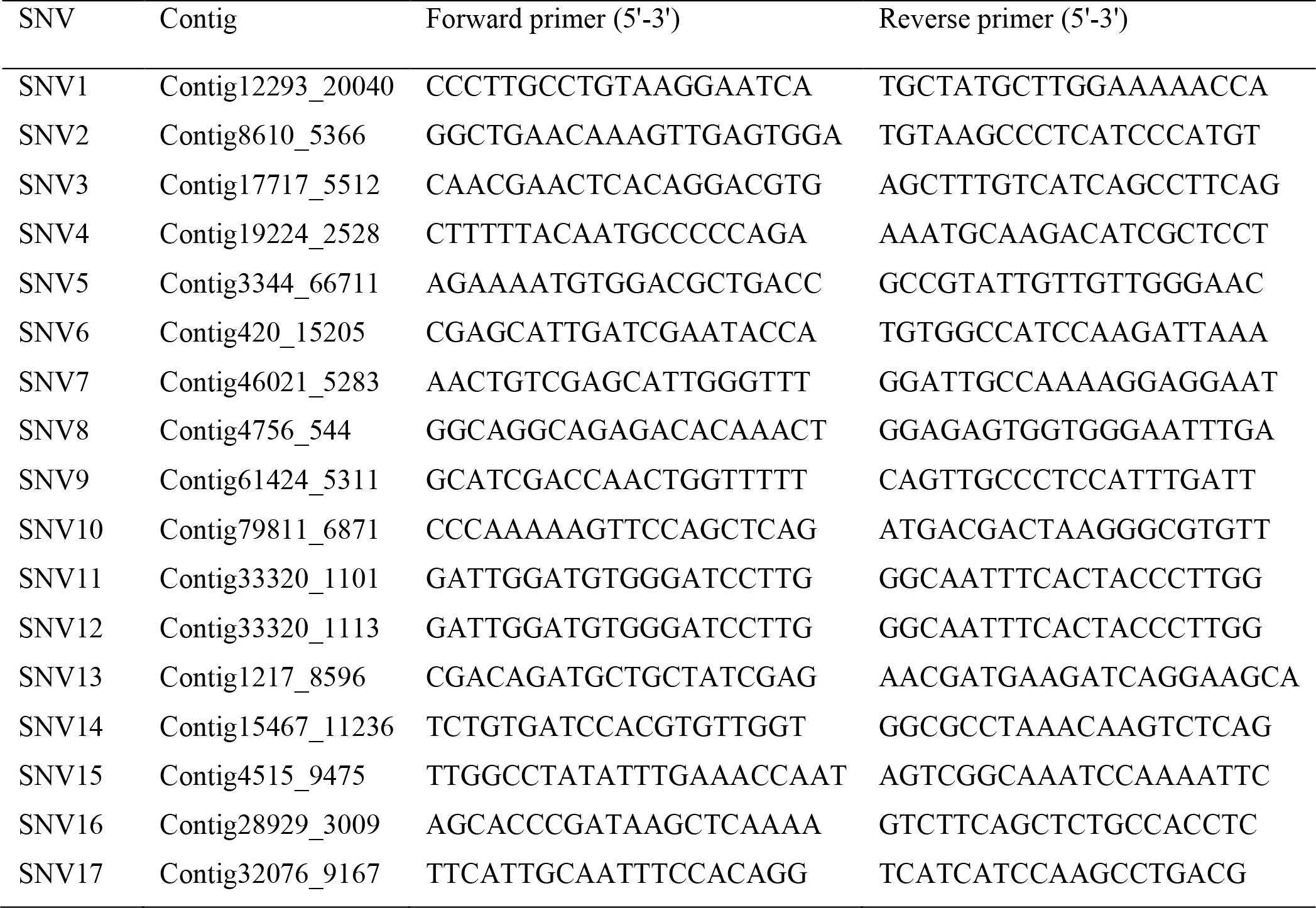
List of primers used for genotyping and Sanger sequencing

## Low rate of somatic mutations in a long-lived oak tree

Sarkar *et al.*

## Supplementary Methods

### Genome assembly

For sample 0, a paired-end library generated 2 × 151,194,704 reads (coverage 40X) and a mate-pair library generated 2 × 107,264,298 reads (coverage 29X). For sample 66, a paired-end library generated 2 × 158,505,474 reads (coverage 42X) and a mate-pair library generated 2 × 124,076,608 reads (coverage 33X). These reads were filtered and trimmed prior assembly using Trimmomatic (v0.3; leading:3, trailing:3, slidingwindow:4:15, minlen:36, custom adapter library)^23^ and assembled using SOAPdenovo2 (v2.04.240, kmer 49)^24^. In a second step the assembly was scaffolded with mate-pairs using the same program. The assembly was further scaffolded with long single-molecule PacBio reads (22 SMRT cells, XL-C2 and P4-C2 chemistry, coverage 19X) and the program AHA (http://www.pacb.com/products-and-services/analytical-software/smrt-analysis/; SMRTPipe 2.0.1 manually driven, settings (5,2,50,70), no gap-filling). Assembled sequences <1000 bp were removed to facilitate further analysis. The genome was extended with all paired-end libraries and SSPACE^25^ (v2.0, -x = 1,z = 0,-k = 5,-a = 0.7,-n = 15,-T = 20,-p = 0,-o = 20,-t = 0,-m = 32,-r = 0.9) and gaps were filled using Gapfiller (v1.10, all paired-end libraries)^26^.

We screened the paired-end libraries for potential non-oak sequences using metaphlan (v1.7.7)^27^. Based on metaphlan results, reference genomes were obtained for the non-oak genomes and the oak scaffolds were filtered against these using blast (ncbi-blast v2.28, >90% sequence identity and E-value <1e-5). The genome was next scaffolded again using the PacBio reads and PBJelly (v14.1.14)^28^. If not further specified, programs were used with their standard settings.

### Gene prediction and annotation

Repetitive elements were analysed by first generating a specific repeat model using RepeatModeler (http://www.repeatmasker.org; v1.0.7, -engine wublast). Repetitive regions in the genome were subsequently masked with the obtained model using RepeatMasker (http://www.repeatmasker.org; v4.0.3). Genes were predicted by generating a *Q. robur* specific gene prediction model for Augustus (v3.0.1)^29^, as described in Tran et al.^30^. Instead of RNAseq reads, we used the UniProtKB reference proteome of *Glycine max* mapped with the splice aware mapper exonerate (V2.2.0, model protein2genome, geneseed 250 –minintron 20, -maxintron 20000)^31^. Using this model we predicted genes and subsequently their encoded proteins for the hard-masked version of the genome (settings: no hints, no UTR predicted, no alternative transcripts). Non-coding elements were annotated using RFAM (v1.5; infernal 1.0.2; blast 2.2.26; hmmer 3.1b1)^32^ in the genome with coding regions masked but repetitive elements unmasked. The predicted proteome was annotated based on homology using the FASTA toolkit (http://www.ebi.ac.uk/Tools/sss/fasta/; v36.3.5e) as following: proteins from the *Glycine max* proteome were first mapped with ggsearch (-b 1 -d 0 -E 1e-5 -m 8 -T 10); proteins that did not map were mapped in a next step with glsearch (-b 1 -d 0 -E 1e-5 -m 8 -T 10) and finally the rest with ssearch (-b 1 -d 0 -E 1e-5 -m 8 -T 10). The functional protein annotation was overtaken from *Glycine max*. For proteins with unknown function in *Glycine max*, we extended the annotation using the OMA database (www.omabrowser.org) and orthologous proteins from *Arabidopsis*. PFAM^33^ was used additionally to obtain functional domain annotations for the proteome and the concatenated proteome annotation was transferred onto the oak genome.

### PCR-seq

A modification of the published RT-PCR-seq method^34^ was used. Briefly, pairs of primers for 50-150 bp amplicons containing the targeted sequence were designed using Primer3. Touchdown PCR amplification was performed in a final volume of 12.5 ml with JumpStart REDTaq ReadyMix (Sigma-Aldrich), a primer concentration of 0.4 mM and 2 ng of gDNA per reaction in 384-well plates. Equal volumes of PCR products were pooled for each DNA template (sample 0 and 66). One ml of each pool was then purified with the QIAquick PCR Purification Kit (Qiagen) following the manufacturer’s instructions. The KAPA LTP Library Preparation Kit (Kapa Biosystems) was used, starting with 500 ng of purified PCR products, to create a library compatible with an Illumina sequencing platform. Clean-ups between enzymatic steps were performed with Nucleospin PCR Clean-up columns (Macherey-Nagel). After ligation of pentabase adapters, libraries were run on a 2 % agarose gel and extracted using the MinElute Gel Extraction kit (Qiagen). Libraries were sequenced on HiSeq 2000 after six cycles of amplification (Lausanne Genomic Technologies Facility). Amplicon reads were aligned, with no mismatches allowed, to a compendium of the expected amplimers that bore the reference allele, the alternate allele identified in the heterozygote sample, as well as the remaining two nucleotides at the variable position; this allowed an unbiased estimation of the error rate generated by the sequencing itself. As this method might have missed *bona fide* changes between the two sampled branches that present other heterozygous sites close by, we also aligned amplicon sequencing reads directly to the reference genome, with mismatches allowed.

### Estimation of the possible missed SNVs

About half of the putative variable sites with confidence scores ≥200 were assessed experimentally by PCR-seq (1,536 out of 3,488 sites). Given the confidence scores of the tested sites, we estimated that we missed fewer than 6 SNVs in the sites not evaluated by PCR-seq. We then evaluated the number of true positives missed within candidates with confidence scores <200 in the list of sites that were heterozygous in only one sample. We fitted a mixture of two distributions, including a normal distribution that should fit the correct calls, modelled on the 1,832,554 sites that were predicted to be heterozygous in both samples (Supplementary Figure 5a). Applying this distribution to the data for sites that are homozygous on sample 0 and heterozygous on sample 66 (case 1, Supplementary Figure 5b), we find that the distribution of correct calls is insignificant compared to the rest. In details, when fitting this normal distribution to the data, the expected number of correct calls with a score < 200 is 5.24. Extrapolating this calculation for sites that are heterozygous on sample 0 and homozygous on sample 66 (case 2), we estimate that we have missed fewer than 11 true SNVs for both cases. We thus estimate a total of 17 missed SNVs (6 with a score ≥200 and 11 with a score <200). Note that we did not assess the presence of larger somatic changes such as copy number variants, small indels, and transposition events.

### Estimation of the false negative rate

A few recent studies have tried to estimate the false negative rate of SNV calling for large genomes assembled with short read sequences^10,35^. The main method used in those studies was to introduce simulated SNVs into the data, and check how well they were recovered. This is not directly applicable to our approach, as we did not merely perform bioinformatics filtering, but also used a large PCR-seq screen to detect SNVs. The comparison of F2 / F4 genotypes is very promising, but cannot either be applied to our data^10^. It is notable that both approaches, in Drosophila^35^ and in Arabidopsis^10^, found very low rates of false negatives, less than 2%. If this holds true for our data (which has higher mean read coverage), then our false negative rate cannot be so large as to challenge our main conclusions. Using this maximum rate of 2%, we can estimate the number of false negatives (FN) as: 0.02=FN/FN + TP, where TP equals 34 estimated true SNVs. This calculation gives 0.69, which suggests that we have only missed ~1 SNV.

### Whole-genome duplication

Simple clustering based on homology, (i.e., clustering the predicted proteins by identity, CD-HIT, min 90% similarity), retrieved 1,098 proteins that have a >90% identity to another protein, which is not suggestive of recent whole genome duplication. Whole genome duplication should lead to an excess of relatively old paralogs, whereas small-scale duplicates are expected to be enriched in very recent paralogs. This can be estimated from the distribution of synonymous distances (dS)^36,37^. We computed the dS on a stringent set of 4,777 paralog pairs with BLAST E-value <1e-10, removing large multigene families (more than 20 members). The distribution of dS values is clearly unimodal, with an excess of low dS values (i.e., young paralogs, Supplementary Figure 6). This also does not support a recent whole genome duplication in the oak lineage.

To address the possibility of a more ancient duplication event, we compared our oak genome reference with itself using “BLAST all versus all” as suggested in Panchy et al.^38^, (i.e., similarity ≥30%, match length ≥150AA and E-value ≤1e-5). Following this procedure we have 49,444 proteins, of which 3,650 are duplicated (7.4%), 2,070 are triplicated (4.1%) and 23.7% are present in more copies with diminishing frequency. In summary, a total of 17,474 oak proteins out of 49,444 appear to be duplicated (35%), which is less than that reported for closely related species (e.g. *Medicago sativa* has about 50,000 genes of which >75% are duplicated, according to Panchy et al.^38^). We then assessed whether the similarity identified above was local, properties of similar domains, or extended along the entire protein, indicative of duplicated proteins. We found only 973 oak proteins that have duplications extending over their entire lengths. In summary, it is possible that the oak genome underwent duplication, as suggested by Panchy et al.^38^, but this event appears to be rather old, as we have very few (<3%) duplicated genes with very high similarity (>90%) and no second peak in the dS distribution (Supplementary Figure 6). It seems unlikely that such a duplication event should compromise the identification of *bona fide* variants. Note that if the duplication would have hindered the capacity to detect these variants, they would not be found in nested sectors of the tree but rather in all 26 samples assessed.

### Analysis of DNA repair genes

Orthologs between *Arabidopsis*, *Prunus persica* (peach) and *Q. robur* were called using the OMA database^39^. One-to-many orthologs, e.g., between *Arabidopsis* and *Q. robur*, represent duplication in the oak lineage since the divergence from *Arabidopsis*; they are also known as in-paralogs of oak. We classified these in-paralogs according to whether the duplication was shared by *P. persica* and *Q. robur* (i.e., one copy in *Arabidopsis* relative to several copies in both the peach and oak genomes), or whether it was peach-or oak-specific (i.e., one copy in *Arabidopsis* and peach, relative to several copies in oak). The number of duplicates was reported as the number of genes that could be called duplicate (i.e., the number of orthologs between each tree genome and *Arabidopsis*, Supplementary Table 5). We then manually compiled a list of *Arabidopsis* genes involved in DNA repair from SwissProt/UniProtKB annotations (Supplementary Table 6). We then counted specifically the number of duplicates for genes involved in DNA repair and reported this as the number of orthologs associated with this function (Supplementary Table 3 and 7).

## Supplementary Discussion

We found that G:C→A:T transitions were the most frequent class of SNVs observed in the Napoleon Oak (Supplementary Figure 7). Ultraviolet (UV) light causes G:C→A:T transitions at dipyrimidine sites in plants^40^. Among the 11 G:C→A:T transitions that we observed, seven were in a dipyrimidine context (Supplementary Table 2). In addition, spontaneous deamination of methylated cytosine leads to thymine change at CpG or CpNG sites [22]. However, there were only three G:C→A:T transitions in such a context (Supplementary Table 2). It thus seems plausible that UV light may have caused most of the G:C→A:T transitions we observed, although other factors, such as cytosine deamination and replication errors, may account for other SNVs. Although the oak lineages sampled have not been separated by any meiosis events, which in yeast was found to elevate the generational mutation rate^41^, they have been exposed to the natural environment, which in *Arabidopsis* is known to significantly enhance mutation rate when compared to a controlled lab environment^42^.

Our results throw new light on explanations proposed for differences in the distribution of mating systems between short-and long-lived plants. While many annuals and short-lived plants have undergone evolutionary transitions from outcrossing to selfing^43^, often involving a loss of self-incompatibility systems^44^, long-lived woody species are more likely to be fully outcrossing^45^, including oaks^46^. Theoretical analysis indicates that a high somatic mutation rate could account for this difference, because somatic mutations would contribute to the genetic load of the population and thus to inbreeding depression, disfavouring self-fertilization^1^. Inbreeding depression is indeed higher in long-lived woody species than annuals^47^, and the observation of higher inbreeding depression caused by within-branch than between-branch selfing points to the accumulation of different deleterious somatic mutations in different sectors of the plant^3^. However, our finding now challenges the notion that the breeding system of long-lived trees is constrained by a high rate of somatic mutations.

The results of our study, in conjunction with those of Burian et al.^14^, have important implications for how we should view one of the most fundamental ways in which plants differ from animals – their absence of a germline. In oak, iterative growth of axillary meristems produces terminal branches that carry stem cells. As in other plants, favourable conditions induce stem cells to produce floral buds and ultimately the gametes of the next generation. These stem cells are functionally analogous to germ cells in metazoans and result from a limited number of divisions that prevent an accumulation of replicative errors.

## References

1. Scofield, D.G. & Schultz, S.T. Mitosis, stature and evolution of plant mating systems: low-Φ and high-Φ plants. Proc. Royal Soc. London B: Biol. Sci. 273, 275–282 (2006).

2. Ally., D., Ritland, K. & Otto, S.P. Aging in a long-lived clonal tree. PLoS Biol. 8, e1000454 (2010).

3. Bobiwash, K., Schultz, S.T. & Schoen, D.J. Somatic deleterious mutation rate in a woody plant: estimation from phenotypic data. Heredity 111, 338–344 (2013).

4. Millet, J. L’architecture des arbres des régions tempérées: son histoire, ses concepts, ses usages. MultiMondes ed., 397 pp (2012).

5. Plomion, C. et al. Decoding the oak genome: public release of sequence data, assembly, annotation and publication strategies. Mol. Ecol. Resour. 16, 254–265 (2016).

6. McKenna, A. et al. The genome analysis toolkit: a MapReduce framework for analyzing next-generation DNA sequencing data. Genome Res. 20, 1297–303 (2010).

7. Iseli, C., Ambrosini, G., Bucher, P. & Jongeneel, C.V. Indexing strategies for rapid searches of short words in genome sequences. PLoS One 2, e579 (2007).

8. Wolfe, K.H., Li, W.H. & Sharp, P.M. Rates of nucleotide substitution vary greatly among plant mitochondrial, chloroplast, and nuclear DNAs. Proc. Natl. Acad. Sci. USA 84, 9054–9058 (1987).

9. Ossowski, S. et al. The rate and molecular spectrum of spontaneous mutations in *Arabidopsis thaliana*. Science 327, 92–94 (2010).

10. Yang, S. et al. Parent–progeny sequencing indicates higher mutation rates in heterozygotes. Nature 523, 463–467 (2015).

11. Scofield, D.G. Medial pith cells per meter in twigs as a proxy for mitotic growth rate (Φ/m) in the apical meristem. Am. J. Bot. 93, 1740–1747 (2006).

12. Romberger, J.A., Hejnowicz, Z. & Hill, J.F. Plant structure: function and development. Springer-Verlag ed., 524 pp (1993).

13. Kwiatkowska, D. Flowering and apical meristem growth dynamics. J. Exp. Bot. 59, 187–201 (2008).

14. Burian, A., Barbier de Reuille, P. & Kuhlemeier, C. Patterns of stem cell divisions contribute to plant longevity. Curr. Biol. 26, 1385–1394 (2016).

15. Edwards, P.B., Wanjura, W.J., Brown, W.V. & Dearn, J.M. Mosaic resistance in plants. Nature 347, 434 (1990).

16. Padovan, A., Lanfear, R., Keszei, A., Foley, W.J. & Külheim, C. Differences in gene expression within a striking phenotypic mosaic *Eucalyptus* tree that varies in susceptibility to herbivory. BMC Plant Biol. 13, 29 (2013).

17. Watson, J.M. et al. Germline replications and somatic mutation accumulation are independent of vegetative life span in *Arabidopsis*. Proc. Natl. Acad. Sci. USA 113, 12226–12231 (2016).

18. Behjati, S. et al. Genome sequencing of normal cells reveals developmental lineages and mutational processes. Nature 513, 422–425 (2014).

19. Lodato, M.A. et al. Somatic mutation in single human neurons tracks developmental and transcriptional history. Science 350, 94–98 (2015).

20. Van der Auwera, G.A. et al. From FastQ data to high-confidence variant calls: the genome analysis toolkit best practices pipeline. Curr. Protocols Bioinfo. 43, 11.10.1–11.10.33 (2013).

21. Myers, E.W. & Miller, W. Optimal alignments in linear space. Comput. Appl. Biosci. 4, 11–17 (1988).

22. Li, H. et al. The Sequence alignment/map (SAM) format and SAMtools. Bioinformatics 25, 2078–9 (2009).

## References

23. Bolger, A.M., Lohse, M. & Usadel, B. Trimmomatic: A flexible trimmer for Illumina Sequence Data. Bioinformatics 30, 2114–2120 (2014).

24. Luo, R. et al. SOAPdenovo2: an empirically improved memory-efficient short-read de novo assembler. GigaScience 1, 18 (2012).

25. Boetzer, M., Henkel, C.V., Jansen, H.J., Butler, D. & Pirovano, W. S. Scaffolding pre-assembled contigs using SSPACE. Bioinformatics 27, 578–579 (2011).

26. Boetzer, M. & Pirovano, W. Toward almost closed genomes with GapFiller. Genome Biol. 13, R56 (2012).

27. Segata, N. et al. Metagenomic microbial community profiling using unique clade specific marker genes. Nature Methods 9, 811–814 (2012).

28. English, A.C. et al. Mind the gap: Upgrading genomes with Pacific Biosciences RS Long-Read Sequencing Technology. PLoS One 7, 11 (2012).

29. Stanke, M., Steinkamp, R., Waack, S. & Morgenstern, B. AUGUSTUS: a web server for gene finding in eukaryotes. Nucl. Acids Res. 32, W309–312 (2004).

30. Tran, V.D. et al. RNA sequencing-based genome reannotation of the dermatophyte *Arthroderma benhamiae* and characterization of its secretome and whole gene expression profile during infection. mSystems 1, e00036–16 (2016).

31. Slater, G.S.C. & Birney, E. Automated generation of heuristics for biological sequence comparison. BMC Bioinformatics 6, 31 (2005).

32. Gardner, P.P. et al. Rfam: updates to the RNA families database. Nucl. Acids Res. 37, D136–140 (2009).

33. Sonnhammer, E.L.L., Eddy, S.R. & Durbin, R. Pfam: a comprehensive database of protein families based on seed alignments. Proteins 28, 405–420 (1997).

34. Howald, C. et al. Combining RT-PCR-seq to catalog all genic elements encoded in the human genome. Genome Res. 22, 1698–1710 (2012).

35. Keightley, P.D., Ness, R.B., Halligan, D.L. & Haddrill, P.R. Estimation of the spontaneous mutation rate per nucleotide site in a *Drosophila melanogaster* full-sib family. Genetics 196, 313–320 (2014).

36. Lynch, M. & Conery, J.S. The evolutionary fate and consequences of duplicated genes. Science 290, 1151–1155 (2000).

37. Vanneste, K., Van de Peer, Y. & Maere, S. Inference of genome duplications from age distributions revisited. Mol. Biol. Evol. 30, 177–190 (2013).

38. Panchy, N., Lehti-Shiu, M. & Shiu, S.-H. Evolution of gene duplication in plants. Plant Physiol. 171, 2294–2316 (2016).

39. Altenhoff, A.M. et al. The OMA orthology database in 2015: function predictions, better plant support, synteny view and other improvements. Nucl. Acids Res. 43, D240–D249 (2015).

40. Britt, A.B. DNA damage and repair in plants. Annu. Rev. Plant Physiol. Plant Mol. Biol. 47, 75–100 (1996).

41. Rattray, A., Santoyo, G., Shafer, B. & Strathern, J. N. Elevated mutation rate during meiosis in *Saccharomyces cerevisiae*. PLoS Genet. 11, e1004910 (2015).

42. Rutter, M.T., Shaw, F.H. & Fenster, C.B. Spontaneous mutation parameters for *Arabidopsis thaliana* measured in the wild. Evolution 64, 1825–1835 (2010).

43. Stebbins, G.L. Variation and evolution in plants. Columbia University Press ed., (1950).

44. Goldberg, E.E. et al. Species selection maintains self-incompatibility. Science 330, 493–495 (2010).

45. Barrett, S.C.H., Harder, L.D. & Worley, A.C. The comparative biology of pollination and mating in flowering plants. Phil. Tran. Royal Soc. London B: Biol. Sci. 351, 1271–1280 (1996).

46. Streiff, R. et al. Pollen dispersal inferred from paternity analysis in a mixed oak stand of *Quercus robur* L. and *Q. petraea* (Matt.) Liebl. Mol. Ecol. 8, 831–841 (2009).

47. Goodwillie, C., Kalisz, S. & Eckert, C.E. The evolutionary enigma of mixed mating systems in plants: occurrence, theoretical explanations, and empirical evidence. Annu. Rev. Ecol. Evo. Syst. 36, 47–79 (2005).

